# Transcriptomic profiling identifies breed-specific immune signatures of Tuberculosis susceptibility in cattle

**DOI:** 10.1101/2024.10.19.619179

**Authors:** Rishi Kumar, Sripratyusha Gandham, Hemant Kumar Maity, Uttam Sarkar, Bappaditya Dey

**Affiliations:** National Institute of Animal Biotechnology, Hyderabad, Telangana, India, PIN 500032; Regional Centre for Biotechnology, Faridabad, Haryana, India, PIN 121001; West Bengal University of Animal and Fishery Sciences. Kolkata, West Bengal, India, PIN 700037

**Author notes:** Corresponding Author: Dr. Bappaditya Dey Scientist-F, National Institute of Animal Biotechnology Hyderabad, Telangana, India - 500032, Email ID. Joint first author (Rishi Kumar, Sripratyusha Gandham).

**Keywords:** Tuberculosis, Bovine tuberculosis, Transcriptome, Susceptibility, PBMC, *Mycobacterium tuberculosis*, *Mycobacterium bovis*, BCG

## Abstract

Tuberculosis (TB) remains a major chronic infectious disease in cattle, particularly challenging in India which hosts the world’s largest and most diverse cattle population farmed in close proximity to its human settlements. This study investigates the hypothesis that genetic and immune variations drive differential TB susceptibility in cattle breeds. Using comprehensive transcriptomic analyses, we examined immune responses in peripheral blood mononuclear cells (PBMCs) from Sahiwal and Sahiwal-Holstein Friesian (SHF) crossbred cattle. Responses to both the virulent *Mycobacterium tuberculosis* and the vaccine strain *M. bovis* BCG were compared. Notably, Sahiwal cattle exhibited a robust early immune response characterized by upregulation of interferon signaling, enhanced cytotoxic activity by CD8+ T and NK cells, and pronounced T cell recruitment and activation pathways compared to SHF crossbreds. PBMCs of this breed also demonstrated superior control of *M. tuberculosis* replication *in vitro*. A distinctive immune signature comprising 8 genes, including *CXCL10, ISG15, CTLA4, SELL, TLR3, MYD88, IRF1, and EOMES* was significantly upregulated in Sahiwal cattle PBMCs, potentially driving their reduced TB susceptibility. These findings underscore the importance of breed-specific immune profiling in devising effective TB control strategies, and could lead to targeted interventions that leverage genetic and immunological insights to mitigate TB in regions with high cattle diversity.

## Introduction

Tuberculosis (TB) in cattle remains a major challenge in livestock farming worldwide, particularly in India, which has the largest and most diverse cattle population (1,2). This population is primarily made up of both indigenous breeds, such as Sahiwal, Gir, Red Sindhi, Tharparkar, Kankrej, and Ongole etc and crossbred varieties that result from breeding with European donor breeds like Holstein Friesian, Jersey, and Brown Swiss (3). While cross- breeding has been widely adopted in India to enhance milk yield, it has also introduced new challenges, including increased susceptibility to regional environmental stress conditions and infectious diseases like TB (4–6). TB in cattle can be caused by various Mycobacterium tuberculosis complex (MTBC) species, including *M. bovis, M. tuberculosis, M. caprae,* and *M. orygis*, each with different levels of infectivity (7). Although *M. bovis* is traditionally considered the primary cause of TB in the bovine, recent studies have highlighted a rising incidence of *M. orygis* in Southeast Asia making the understanding of *Mycobacterium* species- and cattle breed- specific disease resistance increasingly crucial (8–10). Further, zoonotic TB continued to be a public health concern in India, where close human-cattle interactions facilitate cross-species transmission of MTBC species such as *M. tuberculosis* in cattle and *M. bovis* in human (11). In addition, infected cattle, especially those harboring drug-resistant TB strains, pose a serious risk of transmission to humans (12).

The cross-breeding of Indian cattle with exotic breeds has been practiced for over half a century to improve milk production (13). These breeding programs have focused on integrating exotic (mainly European) genetic traits into local cattle populations to boost productivity. Crossbreeds like the Sahiwal-Holstein Friesian (SHF) have become integral to the Indian dairy industry, contributing significantly to the country’s milk supply. According to recent livestock surveys, crossbred cattle account for more than one-third of India’s total cattle population and produce nearly 48% of the nation’s cow milk (4). However, while crossbreeding has succeeded in enhancing milk yield, it has also inadvertently compromised the local environmental adaptability and disease resistance that indigenous breeds naturally possess. Several studies have documented a higher prevalence of TB in crossbred cattle compared to indigenous breeds. For example, research by Thakur et al. reported significantly higher TB positivity in Jersey crossbreds compared to indigenous cattle, while Das et al. found that TB incidence in exotic and crossbred cattle (34.6%) was more than three times higher than in indigenous breeds (10.5%) (14,15). Similar trends were observed in other parts of the world, such as Ethiopia, where Holstein cattle were found to suffer from more severe TB pathology compared to native Zebu breeds (16,17). The trade-off between productivity and disease resistance poses a significant challenge for the livestock farming and industry, as it underscores the need for more nuanced breeding programs that balance both traits.

Our previous study demonstrated that indigenous Sahiwal breed of cattle mounts stronger immune responses and greater control over Mycobacterial growth compared to crossbred SHF cattle (18). Using *M. bovis* BCG vaccine strain and the *M. tuberculosis* H37Ra strain (less virulent), we previously showed that Sahiwal cattle induced higher levels of interferon-gamma (IFN-γ), Interleukin – 17 (IL-17), and nitric oxide, which are crucial for controlling mycobacterial infections (18). Building on this foundation, the current study explores these breed-specific immune responses further by incorporating more virulent strains, such as *M. tuberculosis* H37Rv, along with the *M. bovis* BCG vaccine strain, to conduct comparative transcriptomic analyses. Our aim was to understand the genetic and immunological mechanisms that underlie the differential susceptibility to TB between indigenous and crossbred cattle. Specifically, we analyzed the peripheral blood mononuclear cells (PBMCs) from healthy Sahiwal and SHF crossbred cattle to identify key immune signatures that drive breed-specific disease outcomes. Our results show that Sahiwal cattle mount a robust early immune response, characterized by upregulation of interferon signaling, Toll-like receptor & chemokine mediated signaling, enhanced cytotoxic activity by CD8+ T and NK cells, and superior control of *M. tuberculosis* replication, in contrast to the more subdued responses observed in SHF crossbreds. In addition, we identified a transcriptional signature of 8 genes playing critical role in immune cell recruitment, activation, apoptosis, and inflammation as key mediators of host response to *M. tuberculosis* infection in cattle.

This study highlights the importance of breed-specific immune responses for the development of effective TB control strategies. By identifying the genetic and immunological factors that confer TB resistance and/or susceptibility in cattle, we can inform breeding programs and design targeted interventions that prioritize both productivity and disease resilience. This study represents an important step toward revealing the complex interplay between genetics, immune responses, and disease susceptibility in cattle, with significant implications for improving TB control in regions with diverse cattle populations.

## Materials and Methods

### Animals

Cattle herds from two nearby organized dairy farms in the Nadia area of West Bengal, India, were chosen for this study. Both farms raised a mixed population of crossbred Sahiwal and SHF cattle, and follow similar feeding and management techniques. We took into consideration cows (cattle) those were older than two years, not pregnant throughout the study period, and had tested negative for Tuberculin Skin Test and Myco-PCR negative during the preceding year (18).

### *In vitro* assessment of bovine PBMCs and mycobacterial growth

To comparatively evaluate the growth of virulent mycobacteria within PBMCs from two cattle breeds, an *in vitro* assessment method was employed as reported earlier (18). In addition to the use of *M. bovis* BCG: pMSP12- mCherry fluorescent strain, in this study, a reporter strain of *Mycobacterium tuberculosis* H37Rv was generated using the episomal plasmid pTEC27-Hyg (a gift from Lalita Ramakrishnan, Addgene plasmid #30182; http://n2t.net/addgene:30182). The strain was cultivated to the mid-logarithmic phase in Middlebrook 7H9 medium. Glycerol stocks of the strain were prepared and stored at -80°C following established protocols (19). **Supplementary Table-S1** lists all the plasmids and Mycobacterial strains used in the study. Initially, the fluorescence of the *M. tuberculosis* H37Rv strain was evaluated by growing it in 7H9 medium and monitoring fluorescence levels over a period of 7 days. Subsequently, an in vitro infection was conducted using bovine macrophage (BoMac) cells. For the infection, BOMAC cells were infected with a fresh bacterial culture, grown to mid-logarithmic phase, washed thoroughly with 1X PBS, and resuspended in cell culture media at pre-calibrated dilutions, as previously described (18). The growth of *M. tuberculosis* H37Rv within BoMac cells was monitored every 24 hours over a period of 7 days. Following the establishment of fluorescent mycobacteria and the determination of the multiplicity of infection (MOI) using BoMac cells, bovine peripheral blood mononuclear cells (PBMCs) were isolated from healthy, SITT-negative, and myco-PCR-negative Sahiwal and SxHF cattle according to previously described protocols (18). PBMCs were seeded at a density of 5×10^4^ cells per well into 96-well tissue culture plates for the mycobacterial growth assay. The cells were subsequently infected with mid-logarithmic phase cultures of fluorescent reporter *M. tuberculosis* H37Rv-pTEC27 and *M. bovis* BCG- pMSP12::mCherry at a predetermined MOI of 1:10 (18). Using a fluorescence multimode plate reader, the increase in fluorescence intensity, indicative of mycobacterial growth was assessed on alternative days over the course of seven days.

### RNA extraction

The bPBMCs from both breeds of cattle (n = 3 / group) were seeded into a TC-treated 6-well plate at a density of 1x10^6^ cells per mL. These cells were then infected with *M. bovis* BCG and *M. tuberculosis* H37Rv at a predetermined MOI of 1:10, respectively. Twenty-four hours post infection, total RNA was extracted from bovine PBMCs using the RNeasy Plus Kit (Qiagen Inc, CA, USA) following the manufacturer’s instructions. Any residual genomic DNA was eliminated through additional treatment with RNase-free DNase (Qiagen Inc, CA, USA). Total RNA was eluted in 30μl of RNase free water and stored at -80°C. Subsequently, the quality and quantity of the extracted RNA were assessed using a Nanodrop Spectrophotometer (Nanodrop1000, Thermo Fisher). and the RNA-seq was outsourced to Neuberg Center for Genomic Medicine, Ahmedabad, Gujarat, India.

### Whole transcriptome sequencing and Analysis

RNA quantity check: The quantity of extracted RNA was assessed using the Qubit fluorometer (Thermo Fisher #Q33238) with the RNA HS assay kit (Thermo Fisher #Q32851) as per the manufacturer’s instructions. Additionally, to evaluate purity of extraction, sample concentrations were measured on the Nanodrop 1000. Subsequently, RIN values were determined for the samples using the TapeStation 4150 with HS RNA screen tape. The samples which had the optimum concentration, A260/280 and A260/230 ratios were proceeded for library preparation after one round of purification.

Library preparation and QC: Library preparation was carried out using the TruSeq® Stranded Total RNA kit (Illumina #15032611). Final library quantification was performed with the Qubit 4.0 fluorometer (Thermo Fisher #Q33238) using the DNA HS assay kit (Thermo Fisher #Q32851) according to the manufacturer’s instructions. To determine the insert size of the library, it was analyzed on the TapeStation 4150 (Agilent) with high-sensitive D1000 screentapes (Agilent #5067-5582) following the manufacturer’s protocol. The Quality of the raw fastq reads of the sample was assessed using FastQC v.0.11.9 with the default parameters (20). The raw FASTQ reads underwent preprocessing using Fastp v.0.20.1 (21), followed by quality reassessment via FastQC and summarization using MultiQC (22).

### Mapping of the processed sequencing reads to the reference genome and analysis

To enhance the specificity of alignment, processed Fastp reads were then aligned to the Silva database to filter out rRNA reads using Bowtie2 (23), 2.4.5 (with parameters: --al-conc-gz, --un-conc-gz). The remaining non-rRNA reads were aligned to the STAR indexed Bos taurus genome (source: ENSEMBL; ARS-UCD1.2.107) using STAR aligner v 2.7.9a (24) (with parameters: --outSAMtype BAM SortedByCoordinate, --outSAMunmapped Within, --quantMode TranscriptomeSAM, --outSAMattributes Standard --outFilterScoreMinOverLread 0.33 -- outFilterMatchNminOverLread 0.33). rRNA features were eliminated from the GTF file of Bos taurus. The alignment files (sorted BAM) from individual samples were quantified using featureCounts v. 0.46.1 (25) based on the rRNA-filtered GTF file to obtain gene counts. These gene counts served as inputs for DESeq2 (26) for differential expression estimation with specific parameters: statistical significance threshold --alpha 0.05; and the Benjamini-Hochberg BH for p-value adjustment method:). The ’regularized log’ transformation in DESeq2 was applied for principal component and clustering analysis.

### Analysis of differentially expressed genes (DEGs)

To estimate the differential gene expression between Sahiwal and SHF cattle, subjected to different treatment conditions, the transcript count tables from each treatment were used as an input in the DeSeq2 package in the R software. The gene ids from the count file which did not fulfil the criteria of adjusted p value <0.05 and Log_2_FC ∓ 1 were excluded from further analysis. Venn diagrams were produced using Venny 2.1, Heatmaps were generated following hierarchical clustering, and discriminating variables between comparison groups were identified using a false discovery rate of p < 0.005 or q < 0.2. Functional enrichment analysis was executed employing the g:profiler software (27), to identify the overrepresented biological terms, such as Gene Ontology (GO) categories, biological process.

### Protein-protein interaction (PPI) network analysis

The STRING (Search Tool for the Retrieval of Interacting Genes/Proteins) database (https://string-db.org/) is one of the most comprehensive resources for investigating protein-protein interactions (28). In this study, we submitted both up-regulated and down-regulated differentially expressed genes (DEGs) related to *Mycobacterium tuberculosis* infection from the SHN versus CBN comparison to the STRING database for protein-protein interaction (PPI) network prediction. With a medium confidence level set at 0.4 and default parameters, the entire spectrum of string network types including both physical and functional association was adopted. The network dynamics of the generated gene clusters were then examined by importing the created network into Cytoscape (version 3.10.1). Using the MCODE clustering algorithm, highly connected clusters were identified during network analysis, aiding in the detection of areas within the network that exhibit high connectivity (29). The cluster in the network with the highest MCODE score was chosen for further analysis. Furthermore, the highly connected genes within a network, referred to as the Hub genes were identified by using seven distinct topological analysis techniques MCC, degree, closeness, radiality, betweenness, stress, and maximum neighborhood component (MNC) to identify the key regulators involved in regulating multiple pathways and processes, and they can be critical for understanding the core functions of the network. The key hub genes were then determined using the maximal clique centrality (MCC) technique based on the MCC Score. Additionally, the biological process of genes sets were also analyzed by importing them into CluGo, a cytoscape plugin that integrates data from multiple biological databases, to help interpret the biological significance and functional enrichment of gene lists by mapping them to different functional categories and related pathways (30).

### qRT-PCR for validation of transcriptome data and potential biomarker for tolerance to TB in Sahiwal cattle

cDNA was synthesized from RNA with the Prime script 1st-strand cDNA synthesis kit (Takara) using a combination of random hexamer and oligo dT primers, in accordance with the manufacturer’s instructions. Primer-BLAST (NCBI) was used to create primers for the bovine gene targets (CXCL10, ISG15, CTLA4, SELL, TLR3, MYD88, IRF1, and EOMES), and a CFX96 Touch System (Biorad) was used for real-time PCR. Primers sequences were listed in the **Supplementary Table-S2**. A two-minute initial denaturation and enzyme activation at 95°C was followed by 40 cycles of denaturation at 95°C for 15 seconds. Annealing and extension were then performed for one minute at a temperature ranging from 55°C to 65°C (depending on the target gene) in the real-time PCR technique. The samples were heated from 65°C to 95°C in increments of 0.5 for the purpose of performing a melt curve study, and the fluorescence was observed throughout. Relative gene expression of the target genes was calculated using the 2^−ΔΔCT^ method with RPLP0 as an internal control (31).

### Statistical analysis

Using a combination of statistical software and bioinformatics tools, the statistical evaluation for the study was carried out and has been detailed in the figure legends or under the corresponding sub-sections in the Materials and Methods section. Genes with a false discovery rate (FDR) of less than 0.05 and a log_2_ fold change of more than ±1 were deemed significantly differentially expressed. Differential expression analysis was carried out using DESeq2. A principal component analysis (PCA) was undertaken to analyze the variance and clustering of samples. To see the distribution of genes with differential expression and the patterns of expression, heatmaps and volcano plots were created. A hypergeometric test was used to assess the significance of the enriched GO keywords and pathways; a p-value threshold of less than 0.05 was deemed statistically significant. Graphs were made using R (version: R 4.3.2). GraphPad Prism 9 was also used for data analysis and graph generation in selective cases as indicated in the corresponding figure legend. A 95% confidence interval was obtained by applying a significance threshold of p < 0.05 to all statistical tests.

## Results

### Growth of virulent *Mycobacterium tuberculosis* is subdued in the indigenous cattle-PBMCs

To investigate the ability of indigenous Sahiwal and crossbred SHF cattle to control the growth of virulent *Mycobacterium tuberculosis* (*M. tuberculosis*), we employed a previously established ’bovine PBMC-mycobacteria growth assay’ using a fluorescent reporter strain of *M. tuberculosis* H37Rv (18). The *M. tuberculosis* H37Rv and *M. bovis BCG* carried episomal plasmids, pTEC27-Hyg and pMSP12::mCherry expressing tdTomato mCherry fluorescent protein, respectively facilitating fluorescence based bacterial growth monitoring (**Supplementary Table-S1**) (32). The correlation between bacterial numbers and fluorescence intensity was validated through both 7H9 broth and BoMac cell culture assays (**Supplementary Fig. S1**).

PBMCs were isolated from healthy, single intradermal tuberculin test (SITT)-negative and Myco-PCR-negative Sahiwal and SHF cattle, and then infected with a pre-calibrated multiplicity of infection (MOI) of 1:10 (cells to bacteria) using *M. tuberculosis* H37Rv and *M. bovis* BCG fluorescent strains. Fluorescence measurements were taken every alternate day over a 7-day period to monitor bacterial growth. Initially, the fluorescence levels, indicating bacterial load, remained comparable between PBMCs from both breeds. However, by day five, a clear divergence became evident, with significantly lower *M. tuberculosis* growth observed in Sahiwal PBMCs compared to SHF PBMCs by day seven (**Fig. 1A**). As reported previously (18), a similar trend was also observed with *M. bovis* BCG, where the Sahiwal PBMCs exhibited superior control over bacterial growth (**Fig. 1B**). These results suggest that Sahiwal cattle possess a stronger innate ability to restrain the proliferation of virulent *M. tuberculosis* and *M. bovis* BCG in comparison to crossbred SHF cattle. This heightened control in Sahiwal cattle underscores their potential genetic and immunological advantages in resisting mycobacterial infections.

**Fig. 1.**
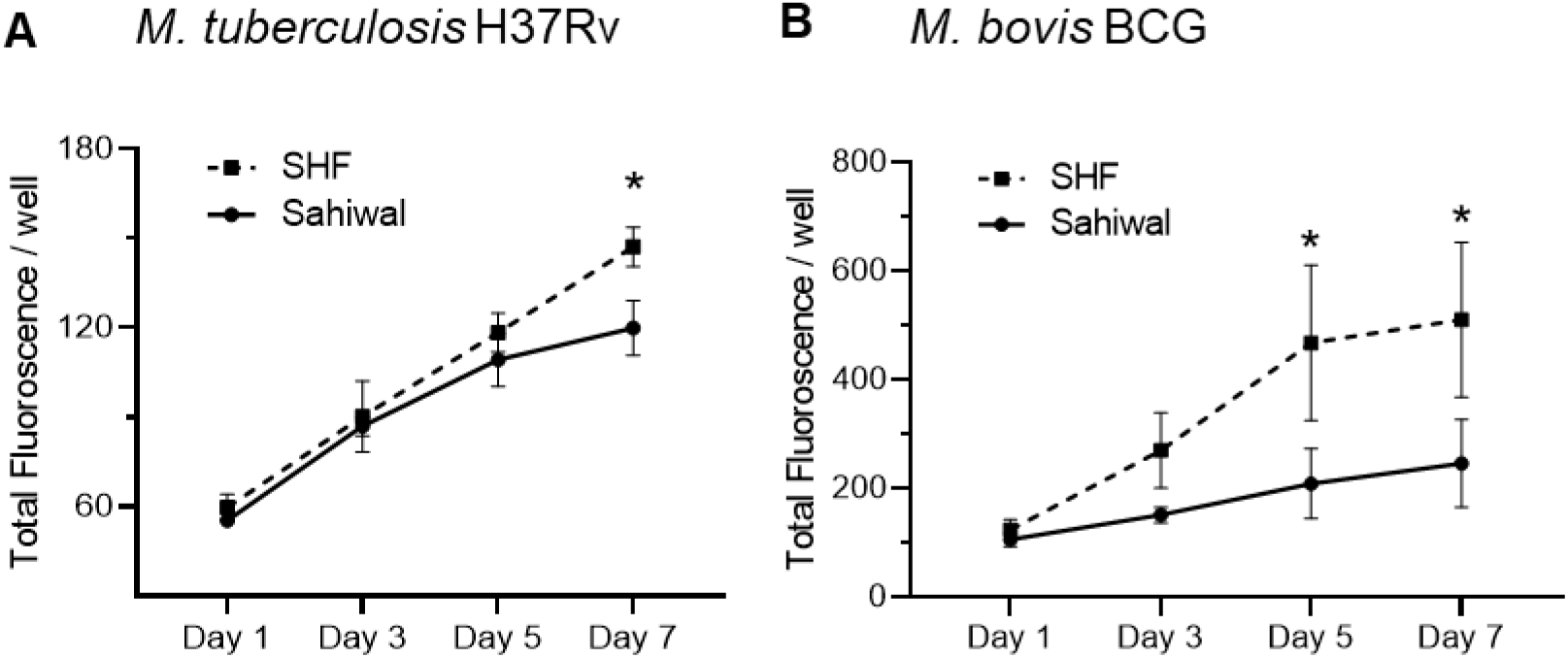
Growth of mycobacteria in bovine peripheral mononuclear cells. PBMCs from Sahiwal and SHF cattle were infected with fluorescence reporter (A) *M. tuberculosis H37Rv-pTEC27;* λ_ex_:554nm and λ_em_:583nm, and (B) *M. bovis* BCG: pMSP12::mCherry; λex:587nm and λem:610nm at an MOI of 1:10 (cell : bacteria). For seven days following infection, the fluorescence intensity of the reporter mycobacteria strain was observed every 48 hours. The data was represented as line graphs by adjoining the mean ± SEM fluorescence / well measured alternative days for each mycobacterial strain. n = 6, *, p<0.05 (t-test). The data is representative of two experiments.

### Significantly altered transcriptome profile in indigenous cattle PBMCs infected with *Mycobacterium tuberculosis* compared to crossbred cattle

To comprehensively understand the innate immune responses during the early phase of *Mycobacterium tuberculosis* infection in cattle and delineate differences between indigenous Sahiwal and crossbred SHF cattle, we performed a detailed transcriptomic analysis of their PBMCs post-infection. The experimental layout of this study is illustrated in **Fig. 2A**. We conducted whole transcriptome sequencing on total RNA extracted from PBMCs of both breeds 24 hours after infection with *M. tuberculosis* and *M. bovis* BCG, and compared the results with uninfected controls. Following sequencing, reads were processed, aligned, and mapped to the *Bos taurus* reference genome ARS-UCD1.2, with normalized read counts generated and deposited in the NCBI GEO database (**GSE277021**). Differential gene expression was analyzed using DESeq2, applying a stringent threshold of a minimum 2-fold change and an adjusted p-value of <0.05 to identify differentially expressed genes (DEGs). The transcriptome analysis workflow is outlined in **Supplementary Fig. S2**.

**Fig. 2.**
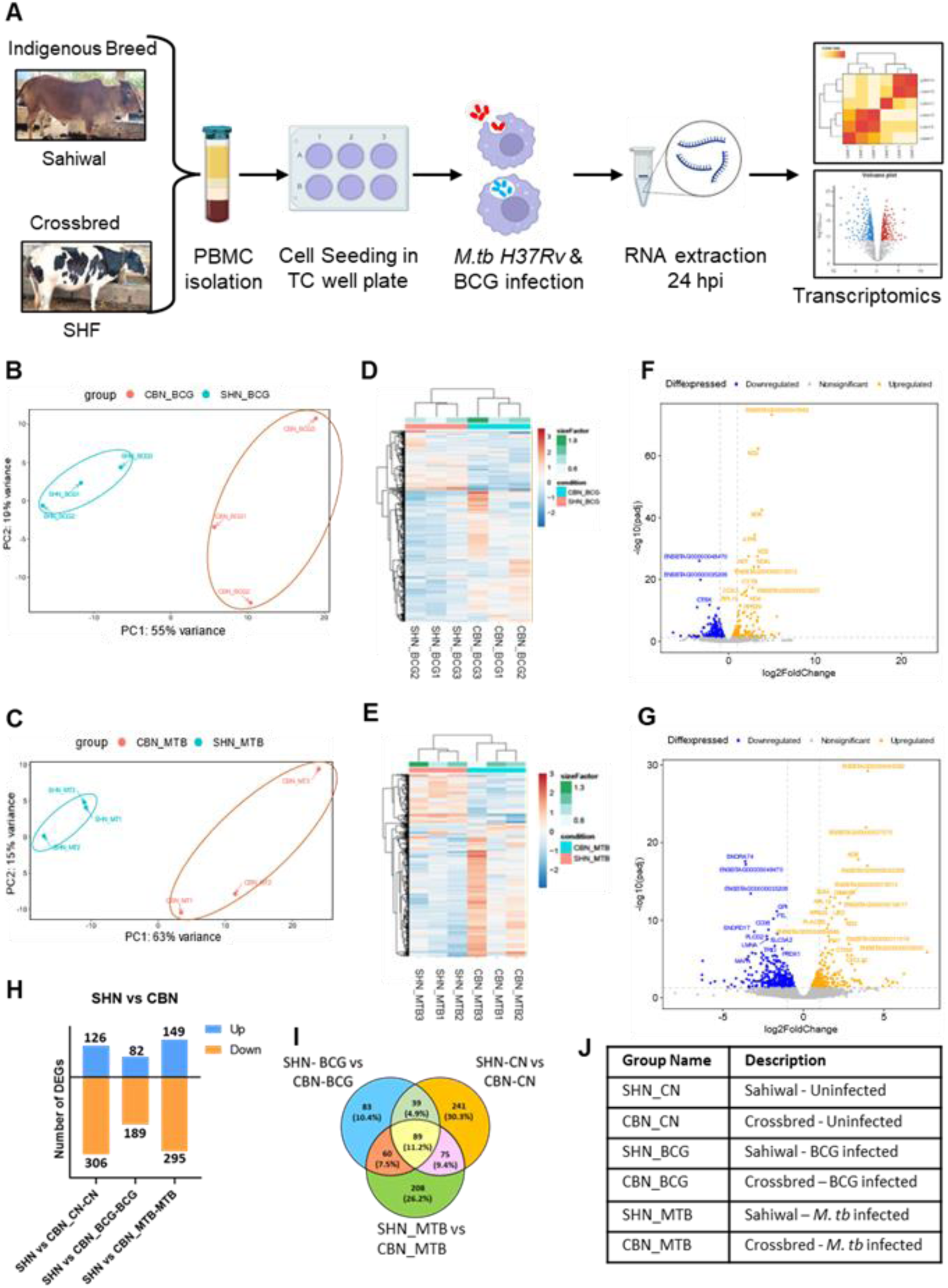
Comparative transcriptome analysis of *M. bovis* BCG and *M. tuberculosis H37Rv* infected PBMCs from Sahiwal and crossbred SHF cattle. PBMCs were isolated from Sahiwal and crossbred cattle and seeded into a 6-well TC-plates. The cells were then infected with *M. bovis* BCG or *M. tuberculosis H37Rv* at an MOI of 1:10, with uninfected PBMCs serving as controls. Total RNA was extracted 24 hours post-infection and subjected to RNA sequencing for transcriptomic analysis. (A). Outline of the experiment; (B, C) The PCA plots representing *M. bovis* BCG, and *M. tuberculosis H37Rv* infection groups- Sahiwal vs. SHF, respectively; (D, E) Heatmap of expression profiles of top 500 genes displaying the relative expression levels across the *M. bovis* BCG and *M. tuberculosis H37Rv* infection groups- Sahiwal vs. SHF, respectively; (F, G) Volcano plot depicting upregulated (orange) and downregulated (blue) differentially expressed genes (DEGs) in *M. bovis* BCG and *M. tuberculosis. H37Rv* infected- Sahiwal vs. SHF with an FDR < 0.05 and Log2FC > 1; (H) Number of Up and Down regulated DEGs under different experimental groups- Sahiwal vs. SHF; (I) Venn diagram-based categorization of DEGs under different experimental groups- Sahiwal vs. SHF; and (J) Description of the experimental group names.

Principal component analysis (PCA) revealed distinct gene expression clusters between Sahiwal (SHN) and crossbred (CBN) PBMCs following infection with *M. bovis* BCG (**Fig. 2B**) and *M. tuberculosis* (**Fig. 2C**). This clear separation indicates significant transcriptional divergence between the two breeds. Hierarchical clustering of the top 500 variable genes reinforced this finding, further validating the differences in immune responses between Sahiwal and crossbred cattle post-infection (**Fig. 2D and 2E**). Volcano plots of the 27,214 differentially expressed genes (DEGs) highlight a significant number of DEGs, with Sahiwal PBMCs showing more robust immune activation compared to crossbred PBMCs when infected with *M. bovis* BCG (**Fig. 2F**) and *M. tuberculosis* (**Fig. 2G**). Interestingly, a similar comparison of uninfected PBMCs between the two breeds also revealed substantial divergence, suggesting that inherent genetic and immune variations between these breeds may influence their baseline immune states and responses to infection (**Supplementary Fig. S3**).

A detailed comparative analysis of differentially expressed genes (DEGs) across the experimental groups revealed marked differences in the transcriptomic profiles of Sahiwal PBMCs compared to crossbred PBMCs. Notably, at baseline, Sahiwal PBMCs exhibited considerable transcriptional variations relative to crossbred PBMCs. Upon infection with both the *M. bovis* BCG vaccine strain and the virulent *M. tuberculosis* strain, these differences were substantiated, with 271 and 444 genes, respectively, showing significant perturbation in Sahiwal PBMCs compared to crossbred PBMCs (**Fig. 2H**). Further analysis identified both unique and shared DEGs within and between the different infection groups, highlighting the distinct immune responses of Sahiwal PBMCs over the crossbred PBMCs. For example, infection with the BCG vaccine strain induced 83 unique DEGs in Sahiwal PBMCs, while infection with *M. tuberculosis* resulted in a much larger set of 208 unique DEGs (**Fig. 3I, Supplementary Data File S1**). The description of the experimental groups was provided in (**Fig. 3J).** These findings underscore the enhanced and distinct immune response capacity of Sahiwal cattle, particularly in response to virulent *M. tuberculosis* infection, compared to crossbred SHF cattle.

**Fig. 3.**
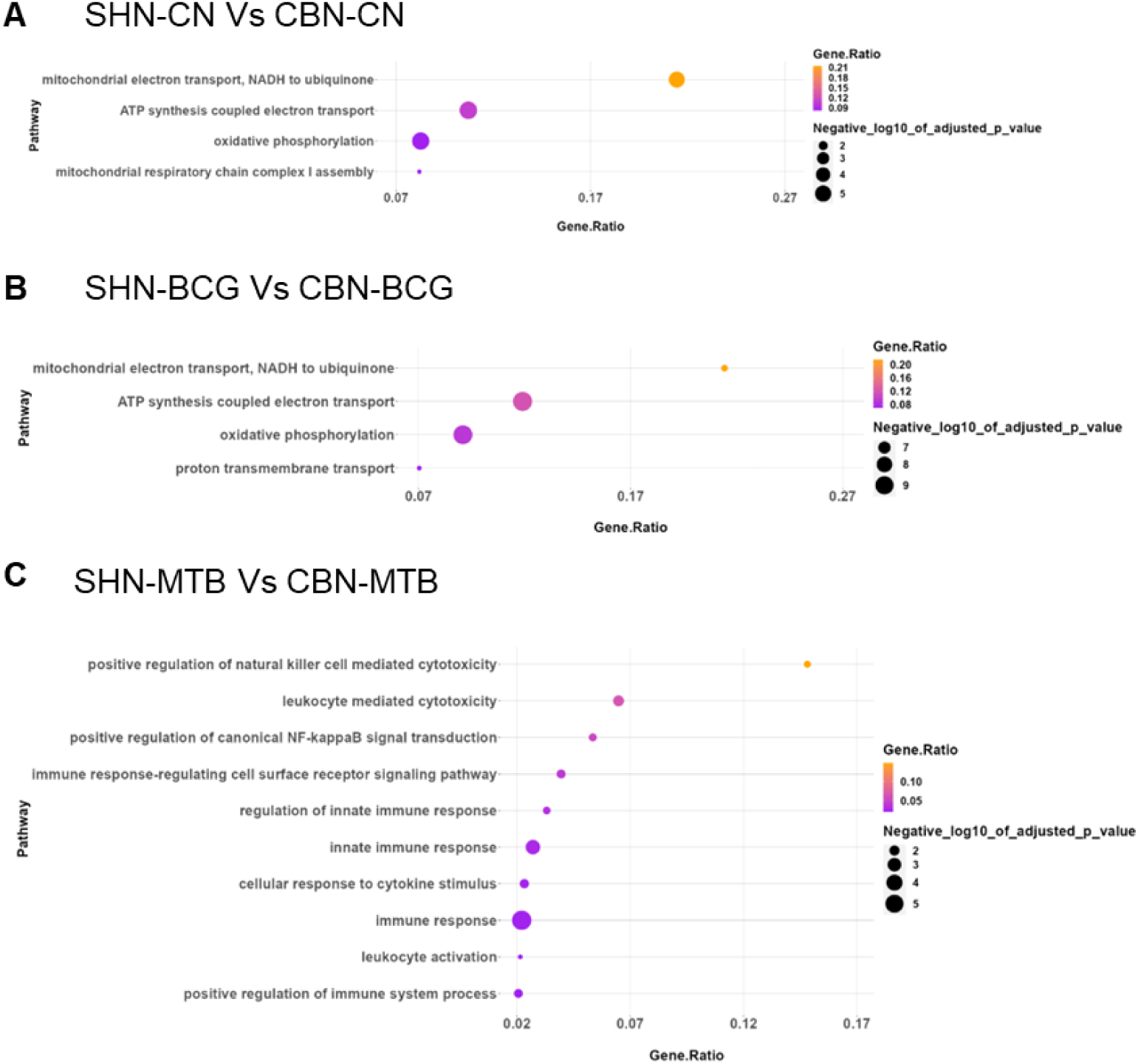
Pathway enrichment analysis of the upregulated genes. Major biological pathway upregulated in case of Sahiwal compared to the crossbred PBMCs, in different experimental conditions: (**A**) Uninfected, (**B**) *M. bovis* BCG infection, and (**C**) *M. tuberculosis* H37Rv infection. The comparison of DEGs was carried out between Sahiwal and crossbred cattle, using crossbred DEGs as the reference. Analysis was conducted using g:Profiler, and figures were generated with R-Studio.

### Functional enrichment analysis of DEGs highlights divergence of biological pathways between Sahiwal and crossbred PBMCs

To elucidate the differences in the biologically and immunologically relevant pathways and processes between the indigenous Sahiwal and crossbred SHF cattle when exposed to *M. bovis* BCG and *M. tuberculosis* infection, we conducted a functional enrichment analysis of unique DEGs belonging to different groups. The functional enrichment via gene-ontology (GO) analysis of the DEGs sourced from uninfected PBMCs of indigenous Sahiwal vs. crossbred cattle revealed a significant enrichment across several biological pathways. Notably, this enrichment highlighted the upregulation of functions including ATP synthesis coupled electron transport, oxidative phosphorylation, thermogenesis, mitochondrial electron transport, and respiratory chain complex in mitochondria (**Fig. 3A)**. It is interesting to note that within the uninfected PBMCs from native breed cattle several pathways are found to be down-regulated, including the toll-like receptor 4 signaling pathway, B cell proliferation, collagen metabolism, interleukin-1 beta production, response to lipopolysaccharide, etc. (**Supplementary Fig. S4A)**. Further, GO analysis of the DEGs in the *M. bovis* BCG infection groups unveiled significant enrichment across various biological processes, such as oxidoreduction-driven active transmembrane transporter activity, oxidative phosphorylation, tricarboxylic acid (TCA) cycle, respiratory electron transport, ATP synthase activity, and structural molecule activity, as illustrated in (**Fig. 3B)**. Moreover, the analysis of downregulated DEGs post-BCG infection in native vs crossbred cattle unveiled regulation of macrophage migration, innate immune system functions, ferroptosis, phagosome activity, and regulation of cell apoptotic processes, as illustrated in (**Supplementary Fig. S4B).** Importantly, the functional enrichment via GO analysis of the upregulated DEGs in Sahiwal vs. SHF PBMCs post-*M. tuberculosis* infection highlights several key biological processes. These include the regulation of innate immune response, immune response-regulating cell surface receptor signaling pathway, positive regulation of canonical NF-kappaB signalling transduction, positive regulation of natural killer cell-mediated cytotoxicity, and innate immune response etc. (**Fig. 3C)**. Functional enrichment analysis of the down- regulated DEGs highlighted several critical pathways including phagocytosis, regulation of programmed cell death, regulation of vesicle-mediated transport, and response to lipopolysaccharide (**Supplementary Fig. S4C)**. These findings shed light on the intricate molecular mechanisms underlying the host’s innate immune response to mycobacterial infection in native Sahiwal breed compared to the SHF crossbred cattle. Excel file of GO analysis for all Up-regulated and down-regulated DEGs is provided in **Supplementary data file S2.**

### Upregulated protein-protein interaction networks (PPIN): Hub genes and functional clusters

The Hub genes are pivotal players within a PPIN, and exhibit extensive connections with other proteins. Recognizing hub proteins is essential for understanding network dynamics and organization and uncovering functional clusters contribute to deciphering the intricate machinery of cells (33). Focusing especially on the *M. tuberculosis* infection specific upregulated DEGs, we constructed protein-protein interactions (PPI) maps using STRING software, and identified the potential interacting networks in the infected PBMCs of indigenous Sahiwal vs. crossbred SHF cattle (**Fig. 4A, and Supplementary data file S3**. After identifying significantly interacting proteins through STRING analysis, we further analyzed upregulated genes in *M. tuberculosis* infected PBMCs using the MCODE algorithm within Cytoscape. Our goal was to pinpoint the fundamental process modulated during early infection with *M. tuberculosis* in the native PBMCs compared to the crossbred cattle PBMCs. Among the upregulated genes a total of three major MCODE clusters were identified encompassing 39 genes, with scores ranging from 3 to 7.31 **(Supplementary data file S4)**. MCODE Cluster-1 encompasses pathways related to chemokine signaling, inflammatory response, and protein synthesis **(Fig. 4B)**. Conversely, Cluster-2 is associated with signaling and transport, protein degradation, IFN-γ signaling, and Type-1 IFN signaling **(Fig. 4C)**. Mucosal immunity and B-cell development are attributed to Cluster-3 **(Fig. 4D)**.

**Fig. 4.**
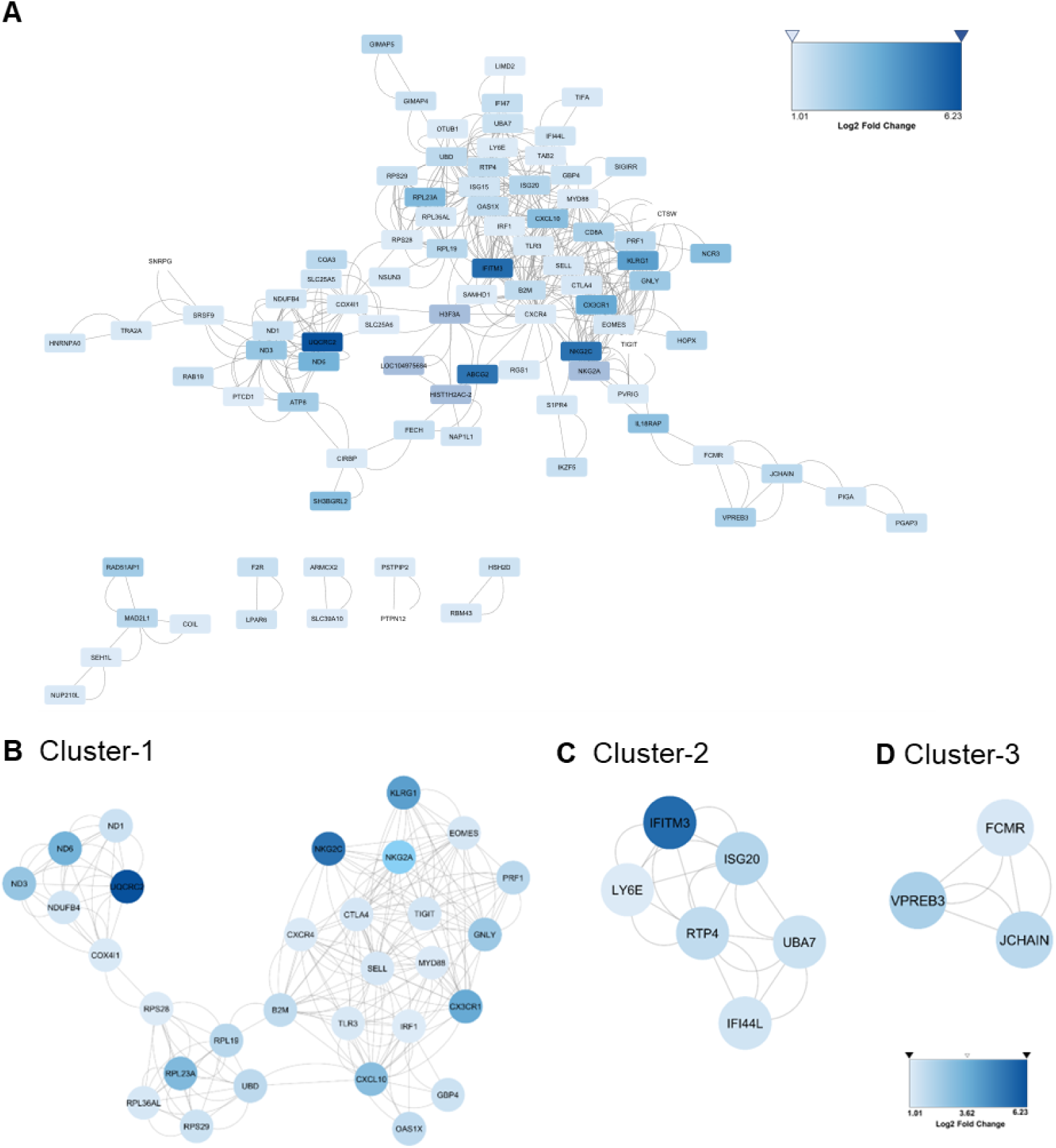
Global and critical gene network analysis of upregulated genes in *M. tuberculosis*-infected groups: Sahiwal vs crossbred. **(A)** Depiction of global networks of all the upregulated genes in case of Sahiwal PBMCs compared to the crossbred cattle via STRING based network analysis; Further, MCODE based analysis in the Cytoscape identified three critical gene clusters: (**B**) Cluster 1: Inflammatory and cellular response, **(C)** Cluster 2: Immune Response, and **(D)** Cluster 3: Humoral Immunity.

Further, to examine the densely interconnected upregulated modules within our dataset, we utilized Cytohubba analysis, a specialized Cytoscape plugin for network analysis (33). Using 7 different features on Cytohubba analysis reveals a total of 34 hubba nodes (**Supplementary Fig. S5 and Supplementary data file S4**). Further ClueGO analysis of these 34 hub-genes reveals three major pathways, namely Interferon gamma production (45.50%), Toll-like receptor signaling pathway (45.50%), and Chemokine mediated signaling pathway (9.09%) that were upregulated during early infection of Sahiwal PBMCs with the virulent *M. tuberculosis* compared to the crossbred SHF cattle (**Fig. 5)**.

**Fig. 5.**
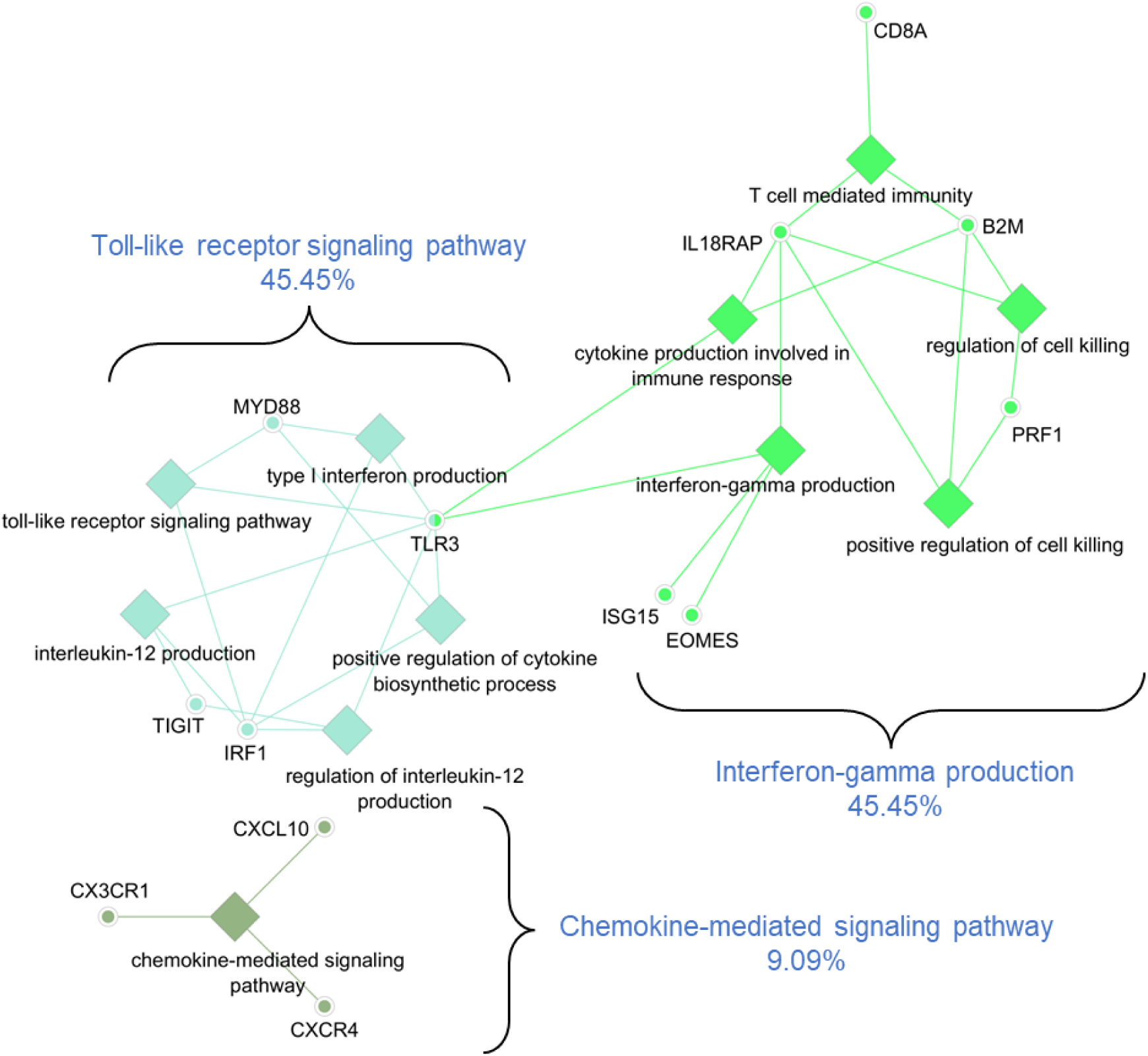
Pathways analysis of the Cytohubba nodes. Signaling network depicting key pathways mapped to the hub-nodes and highly interconnected genes that were upregulated during early infection of Sahiwal PBMCs with the virulent *M. tuberculosis* H37Rv compared to the crossbred SHF cattle. This analysis was performed using the ClueGO plugin in the Cytoscape.

### Transcriptional signature of susceptibility and/or resistance to *M. tuberculosis* infection in cattle

To identify key transcriptional drivers of host immunity against TB in indigenous Sahiwal cattle, we curated a list of 48 DEGs based on the MCODE and CytoHubba analysis (**Fig. 6A, and Supplementary data file S4)**. This initial list was further refined by targeting genes specifically associated with immune responses to *M. tuberculosis* infection, based on information from Uniprot, AmiGo, and InnateDB databases. This led to the identification of 12 critical immune-related genes (*SELL, CTLA4, CXCL10, TLR3, EOMES, ISG15, IRF1, MYD88, IL18RAP, KLRG1, NKG2C,* and *VPREB3*) that play pivotal roles in enhancing host defense against TB.

**Fig. 6.**
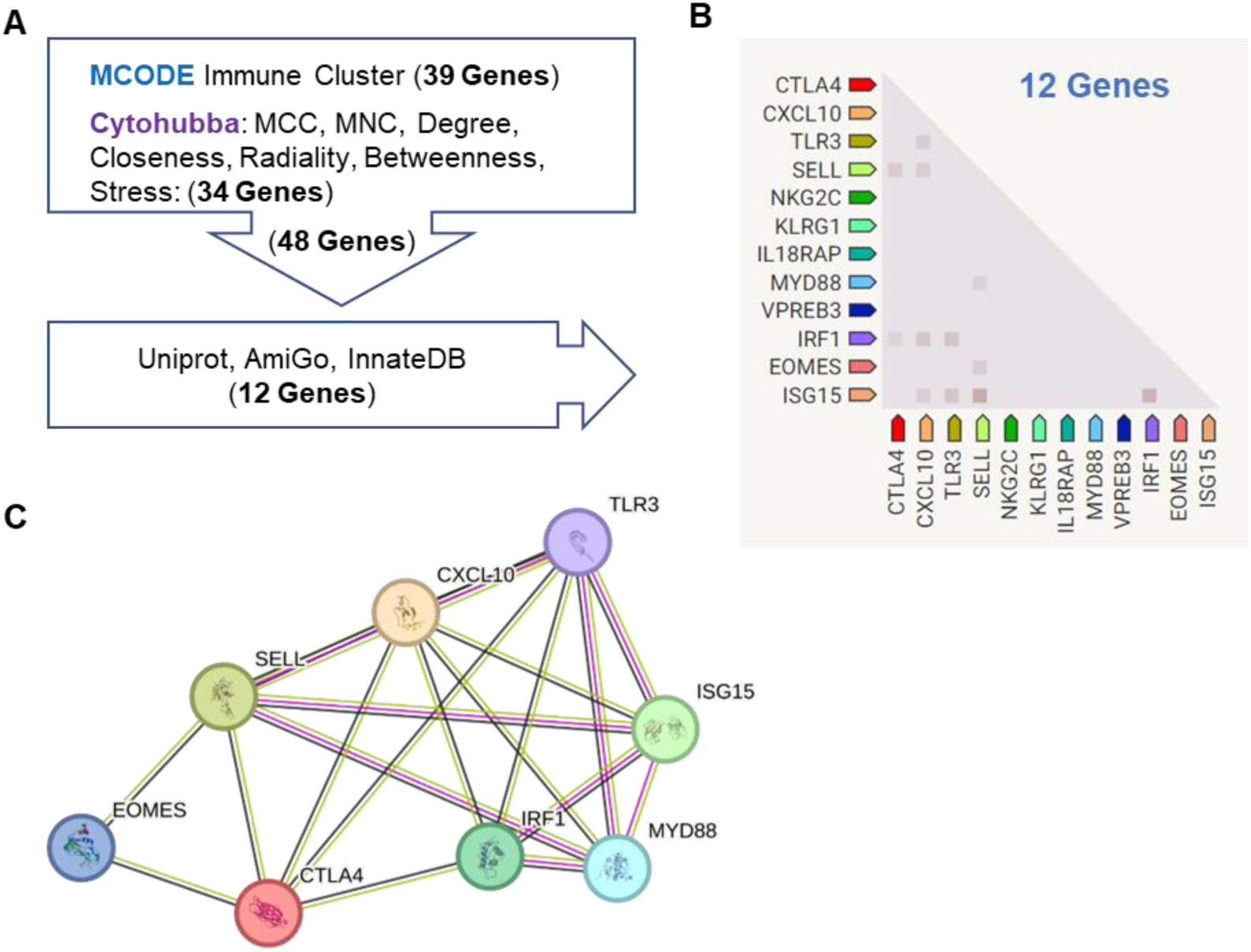
Transcriptional signature of susceptibility and/or resistance to TB in cattle. **(A)** A total of 48 genes were identified via MCODE and CytoHubba analysis and curated via various databases to shortlist 12 genes with known roles in TB immunity; **(B)** Co-expression analysis of 12 shortlisted genes using STRING software; and (**C**) STRING based network of 8 signature genes.

Further co-expression analysis using the STRING database revealed that 8 of these genes (*CXCL10, ISG15, CTLA4, SELL, TLR3, MYD88, IRF1,* and *EOMES*) consistently co-expressed with at least one other gene within the group (**Fig. 6B**, and **Supplementary data file S5**. Network analysis of these 8 genes highlighted their interconnectedness, forming a tightly linked cluster (**Fig. 6C**). The upregulation of this gene cluster in Sahiwal cattle, compared to crossbred SHF cattle, following *M. tuberculosis* exposure suggests that these genes represent a transcriptional signature of robust host immunity to TB in indigenous cattle. The differential regulation of these genes may be a key factor influencing the susceptibility or resistance of different cattle breeds to TB, highlighting their potential as biomarkers for disease resistance in cattle.

### qRT-PCR based validation of RNA-seq data

To validate the transcriptional signature derived from RNA-seq data, we performed qRT-PCR on the 8 selected genes. Our qRT-PCR confirmed the significant upregulation of all the 8 genes - *CXCL10, ISG15, CTLA4, SELL, TLR3, MYD88, IRF1,* and *EOMES* which was in agreement with the RNA-seq data (**Fig. 7**).

**Fig. 7.**
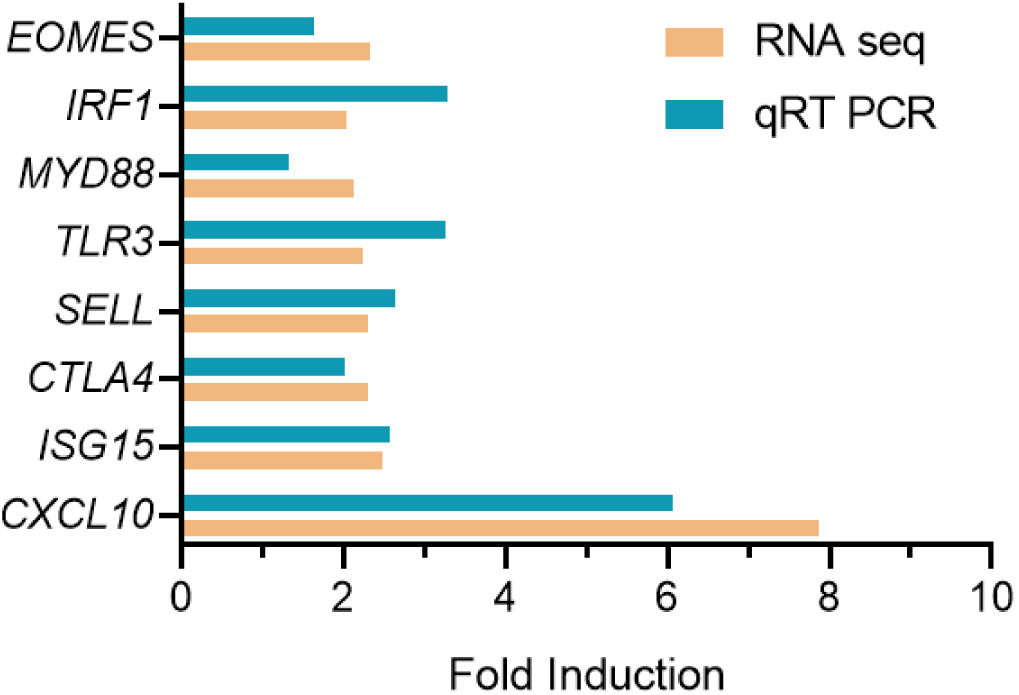
Relative mRNA expression of selected genes by qRT-PCR and RNA seq. Real-time RT-PCR was conducted on 8 genes (*EOMES, ISG15, SELL, IRF1, CTLA4, MYD88, TLR3,* and *CXCL10*). 60S acidic ribosomal protein large (*RPLP0*) was used as the internal control. Fold expression was calculated using the 2^-ΔΔCT^ method. Bars represent the average fold change values from qRT-PCR and RNA-seq analysis, comparing the *M. tuberculosis*-infected PBMCs from indigenous Sahiwal over the crossbred SHF cattle.

## Discussion

Host susceptibility and/ or resistance to TB are shaped by multiple factors, including genetic makeup, nutrition, age, underlying health, and environmental conditions (34). These factors modulate host immune responses, influencing the ability to control or promote the growth of the tubercle bacilli, with a plethora of cellular defense pathways playing key roles (35). Multiple investigations have shown that genetic diversity among organisms significantly influences variations in immune responses to TB (36,37). India, with the largest cattle population globally, has a rich diversity of indigenous breeds that have adapted to various environments, developing resilience and disease resistance. While cross-breeding programs have successfully increased milk production, the introduction of exotic genes has led to genetic dilution, potentially reducing the resilience of these breeds to environmental stress and diseases (6). Crossbred cattle are more susceptible to infections, making health management and disease prevention crucial. Proper nutrition, housing, and veterinary care are essential for maintaining their well-being (4). However, the effects of cross-breeding on susceptibility to infectious diseases like TB, and the underlying genetic and immunological mechanisms, remain underexplored.

In this study, we compared the immune responses of two key dairy cattle breeds in India: the indigenous Sahiwal and the crossbred SHF, which carries 50%-62.5% exotic genetic inheritance. The health status of the animals, including their confirmed mycobacterium-free status, was verified and reported in our previous work (18). Our earlier study provided proof-of-principle that crossbred SHF cattle exhibited a higher susceptibility to Mycobacterium infection compared to the native Sahiwal breed, based on experiments using avirulent *M. bovis* BCG and less virulent *M. tuberculosis* H37Ra strains (18). In the current study, we extended this investigation by assessing the comparative susceptibility of these breeds to the virulent *M. tuberculosis* H37Rv strain. We aimed to explore the relationship between global immune response signatures and the susceptibility or resistance phenotype in these cattle. Using whole transcriptomic analysis, we examined the PBMCs from Sahiwal and SHF cattle 24-hours post-infection with *M. tuberculosis* H37Rv and *M. bovis* BCG strains, providing insights into the breed-specific immune responses driving TB resistance or susceptibility.

Our findings demonstrate that indigenous Sahiwal cattle possess a significantly stronger innate immune response to *M. tuberculosis* infection compared to crossbred SHF cattle. Using a bovine PBMC-mycobacteria growth assay with a fluorescent reporter strain of *M. tuberculosis* H37Rv, we observed a marked divergence in bacterial growth between the two breeds. While the initial bacterial load in PBMCs from both breeds was similar, by day five, Sahiwal PBMCs began to exhibit superior control over *M. tuberculosis* replication, which was significantly more pronounced by day seven. A similar trend was seen with the M. bovis BCG strain, where Sahiwal PBMCs also outperformed SHF PBMCs in restricting bacterial growth. This differential ability to subdue the growth of virulent *M. tuberculosis* suggests that Sahiwal cattle may possess inherent genetic and immunological advantages that contribute to their resistance.

Following our observation that Sahiwal PBMCs were more effective at controlling the growth of *M. tuberculosis* compared to SHF PBMCs, we sought to explore the underlying molecular mechanisms by performing a transcriptomic analysis. The differential gene expression analysis revealed significant transcriptional divergence between the two breeds, further reinforcing the idea that indigenous Sahiwal cattle have a distinct and more robust immune response. PCA and hierarchical clustering of the transcriptomic data showed clear separation between Sahiwal and SHF PBMCs post-infection, with Sahiwal cattle exhibiting a broader and more dynamic range of DEGs following infection with both *M. bovis* BCG and *M. tuberculosis* H37Rv. These findings point to intrinsic genetic and immune differences that likely influence TB resistance. Interestingly, the transcriptomic analysis of uninfected PBMCs also revealed significant differences between the two breeds, indicating that the inherent immune state of the Sahiwal cattle may contribute to their superior ability to combat infection.

Further functional enrichment analysis of the DEGs highlighted key biological pathways that were upregulated in Sahiwal PBMCs post-infection. Pathway analysis of up-regulated DEGs from uninfected Sahiwal vs SHF group identified crucial pathways such as mitochondrial electron transport chain (mtETC), ATP synthesis, oxidative phosphorylation, and mitochondrial respiratory chain complex assembly. Mitochondrial reactive oxygen species (mtROS) and oxidative phosphorylation generated through the mtETC are important innate immune attributes that help macrophages mount antibacterial responses (38). Further, the proton membrane transport pathway is also enriched in BCG infected Sahiwal vs SHF cattle PBMCs. In the immune cells, Ion channels and transporters mediate the transport of divalent cations that have important role as second messengers to regulate intracellular signaling pathways (39). Moreover, uniquely up-regulated DEGs of Sahiwal PBMCs infected with *M. tuberculosis* enriched important innate immune pathways. These included pathways related to innate immune responses, such as NF-kappaB signaling, natural killer cell-mediated cytotoxicity, and cytokine signaling. In particular, the upregulation of genes involved in immune regulation and host defense, such as CXCL10, ISG15, and MYD88, suggests that these pathways play a critical role in the enhanced immune response observed in Sahiwal cattle. This enrichment of immune-related pathways underscores the stronger immune defense mechanisms in Sahiwal cattle, which likely contribute to their greater resistance to TB (40). In contrast, crossbred SHF cattle showed downregulation of several critical immune pathways, including those involved in phagocytosis, cell death regulation, and vesicle-mediated transport. These deficiencies in key immune processes may contribute to their reduced ability to control *M. tuberculosis* infection. The differential regulation of these immune pathways between the two breeds suggests that genetic dilution through crossbreeding may have compromised the immune competence of SHF cattle, making them more vulnerable to TB.

Genes involved in enriched pathway play an important role during mycobacterium infection; development of CD8+ T cell and NK cells, activation of NK cells enhances their ability to kill infected cells and produce cytokines, facilitate adhesion and migration of NK cell, monocytes and T-cell subsets to site of inflammation, activates downstream NF-kB and MAPK pathways that lead to inflammatory responses, enhanced cytotoxic activity of NK cells (41,42). Moreover, presentation of endogenous antigen to CD8+ T cells through MHC class I molecules, enhance the production of IFN-γ in NK cells and T cells, improve the antibacterial defence mechanism and control the intracellular mycobacterium replication (43). Notably, nuclear factor κ-light-chain-enhancer of activated B cells (NF-κB) has been found to play a unique mediator role in providing a pro-inflammatory response to chronic inflammatory disease processes by promoting the activation of macrophages and the release of various cytokines such as IL-1, IL-6, IL-12, and TNF-α (44).

To further pinpoint the molecular drivers of TB resistance in Sahiwal cattle, we identified key transcriptional signatures through protein-protein interaction (PPI) networks via MCODE and hub gene analysis. By focusing on upregulated DEGs specific to *M. tuberculosis* infection, we uncovered four major functional clusters: Chemokine signaling, Toll-like receptor signaling, Type -1 and Type-2 IFN signaling, and B-cell based mucosal immunity, all of which were highly active in Sahiwal PBMCs. These pathways are crucial for coordinating effective immune responses, particularly in controlling bacterial infections.

Chemokine mediated signaling pathway and positive regulation of cytokine biosynthesis process involved in the beginning and execution of the organized and sequential recruitment and activation of cells into *M. tuberculosis*- infected lungs are crucially dependent on chemokines and cytokines (45). Furthermore, Type- I interferons may influence the function of alveolar macrophages and myeloid cells, stimulate abnormal neutrophil extracellular trap responses, inhibit the production of prostaglandin 2, and enhance cytosolic cyclic GMP synthase inflammatory pathways (46). Type- 2 interferons, especially IFN-γ is a crucial cytokine in the immune response against TB (47). When macrophages are activated by IFN-γ, their ability to kill microbes is enhanced, enabling the formation of phagolysosomes that deprive mycobacteria of vital nutrients like iron and expose them to antimicrobial peptides as well as reactive oxygen and nitrogen species, which contribute to bacterial destruction (48). A decrease in IFN- γ production leads to reduced macrophage activity, allowing for increased mycobacterial proliferation (49). Toll-like receptor signaling pathway, which stimulate Th1-biased immune responses, facilitates the development of dendritic cells, and activates antimycobacterial activity crucial for initiating and maintaining the immune response in tuberculosis (50). Moreover, the role of IL-12 in both innate resistance and adaptive immunity is crucial due to its activation of the JAK-STAT signaling pathway. This activation leads to the production of IFN-γ from CD4+ T cells, natural killer (NK) cells, and NKT cells during the initial stages of the immune response. Additionally, IL-12 promotes the differentiation of CD4+ T cells into Th1 effectors, which is important for protecting against bacterial infection (51). The findings indicate that PBMCs derived from native breed cattle exhibited a more robust immune response to Mycobacterium infection in comparison to PBMCs from crossbred cattle.

The identification of 8 key immuno-modulatory genes (*CXCL10, ISG15, CTLA4, SELL, TLR3, MYD88, IRF1,* and *EOMES*) as a potential signature of immunity to TB in cattle highlights their critical role in immune cell recruitment, activation, apoptosis, and inflammation. **Fig. 8** depicts a summary of the signaling networks involving these key mediators of immunity in TB.

**Fig. 8.**
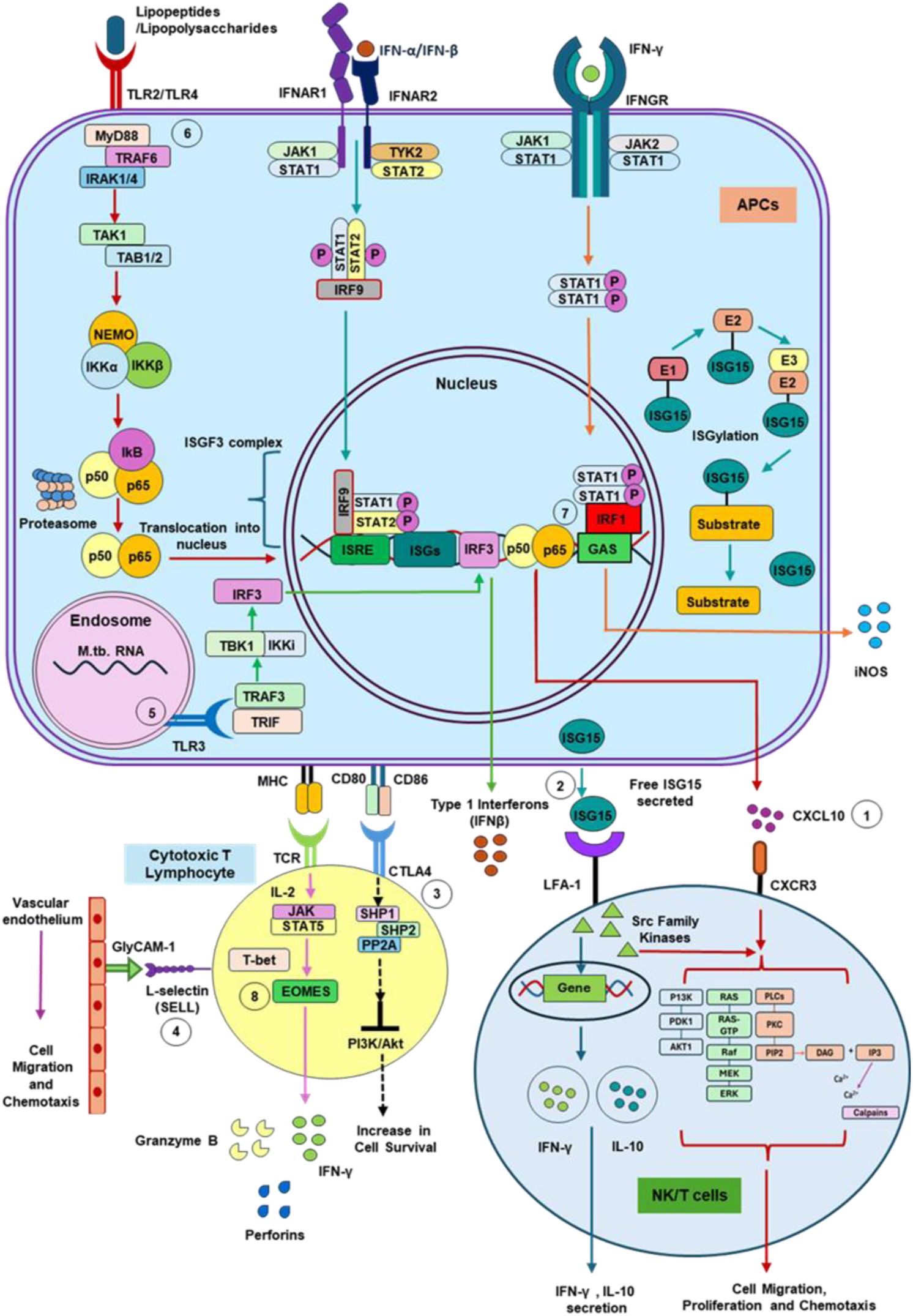
Key molecular players and their signaling networks regulating immunity to TB in cattle. The figure illustrates the key signaling pathways involving the shortlisted immuno mediators during TB. (1) CXCL10 recruits immune cells to infection sites, primarily produced in response to IFN-γ and type I interferons. It binds to CXCR3 on T and NK cells, activating the JAK-STAT and PI3K-Akt pathways, promoting migration and immune activation; (2) ISG15 is a key mediator of Type I interferon response. IFN-α/β are produced during TB upon pathogen recognition by PRRs like TLRs and these bind to the IFNAR, activating the JAK-STAT pathway, leading to the formation of the ISGF3 complex, which induces ISG transcription via ISGylation. ISGylation is a process where ISG15 conjugates to target proteins, stabilizes proteins and boosts key cytokine production, including IFN-γ and IL-10, modulating TH1 and Th2 responses; (3) CTLA-4, an immune checkpoint on activated T cells, downregulates activity during TB by binding to CD80/CD86 on antigen-presenting cells, blocking co-stimulatory signals and initiating inhibitory pathways to prevent excessive immune responses; (4) L-selectin (SELL) facilitates leukocyte migration to infection sites by binding to endothelial glycoproteins like GlyCAM-1, promoting adhesion and transmigration; (5) TLR3, located on endosomes, recognizes double-stranded RNA from *M. tuberculosis*, activating the TRIF pathway to produce pro-inflammatory cytokines via IRF3; (6) MYD88 is an adaptor protein that mediates signaling from most TLRs during *M. tuberculosis* infection, activating the NF-κB and MAPK pathways to enhance pro-inflammatory cytokine production; (7) IRF-1 regulates immune response genes and promotes iNOS production upon IFN-γR signaling, which is crucial for controlling *M. tuberculosis* infection; and (8) EOMES is a transcription factor critical for CD8+ T cell differentiation in TB activating the PI3K-Akt and mTOR pathways. It works with T-bet to promote Th1 polarization and effector cytokine production, enhancing CD8+ T cell cytotoxic functions.

In the context of TB, CXCL10 is a chemokine that directs Th1 cell migration to *M. tuberculosis*-infected tissues, promoting an effective immune response (52). Elevated levels of CXCL10 have been observed in active TB patients, indicating its involvement in the inflammatory response associated with the disease (53). While CXCL10 is vital for immune defense, its dysregulation can lead to excessive inflammation, contributing to tissue damage during TB infection (45). The type 1 IFN inducible protein ISG15 encodes ubiquitin-like protein and mediates ISGylation, which is a potent immunological strategy employed by hosts to combat intracellular infections such as *M. tuberculosis* (54). ISG15 functions as an extracellular and intracellular signaling molecule, impacting the immune response (55). The diverse functions of the ISG15 emphasize its adaptability in exerting influence on an extensive range of cellular pathways, including DNA damage response, autophagy, antiviral response, and processes associated to cancer (56). CTLA-4 is an extensively investigated immunological checkpoint molecule which is produced when T cells are activated and exerts a detrimental effect on its function. CTLA4 regulates T cell response to during TB by serving as an immunological checkpoint, controlling T cell activation, potentially influencing the immune defense capabilities of the immune system (57). L-selectin (SELL), a cellular surface protein, is present on several types of white blood cells, such as T cells and neutrophils. The protein plays a crucial role in enabling the attachment of leukocytes to endothelial cells, therefore enhancing their migration into tissues containing *M. tuberculosis* (58). SELL function is complemented by pattern recognition receptors (PRRs) that recognize *M. tuberculosis* components, triggering signaling pathways that enhance immune responses (59). The interaction between SELL and PRRs promotes the activation of macrophages and T cells, leading to the production of key cytokines like IFN-γ and TNF-α, which are vital for controlling *M. tuberculosis*. infection (59,60).

The Toll-like receptor 3 (TLR3) plays a crucial role in the immune response to *M. tuberculosis* infection by mediating the production of cytokines that influence both innate and adaptive immunity. TLR3 recognizes mycobacterial RNA, leading to the activation of signaling pathways that regulate immune responses (61). TLR3- mediated signaling not only activates innate immune cells but also shapes the adaptive immune response, promoting the differentiation of T cells that are critical for long-term immunity (62). The MyD88 is critical for downstream of all TLR signaling except TLR3 (63). MyD88-deficient mice are susceptible to various pathogens (64), In response to *M. tuberculosis* infection, MyD88-deficient macrophages can express proinflammatory cytokines but are defective in TNF, IL-12, and NO production (65). In a delayed manner, they can also facilitate the expansion of IFN-γ-producing antigen-specific T cells (66). *M. tuberculosis* infection in mice lacking MyD88 is fatal, since MyD88 signaling is essential for generating a sufficient innate and acquired immunological response to *M. tuberculosis* (65).

Interferon regulatory factor 1 (IRF1) enhances macrophage activation, promoting the expression of pro-inflammatory cytokines such as IL-6 and TNF-α, which are crucial for combating *M. tuberculosis* infection (67). IRF-1 is implicated in the development and function of T cells, particularly CD4+ T cells, which are essential for a Th1 cell-mediated immune response against TB (68). Finally, Eomesodermin (EOMES) is involved in the development of cytotoxic T lymphocyte activity via interleukins and other transcription factors (69). EOMES is critical to regulate the expression of key immune molecules such as IFN-γ, perforin, and granzyme B, and regulates the effector T cell response (70). The variable expression of these genes may significantly affect the immune system’s ability to combat or survive TB, therefore influencing the development, intensity, and general susceptibility of the disease.

While this study provides valuable insights into the differential immune responses of indigenous and crossbred cattle to *M. tuberculosis*, there are several limitations that warrant further exploration. One key limitation is the exclusive use of *M. tuberculosis* as the virulent pathogen, without including other relevant pathogens such as *M. bovis* and *M. orygis*. This may limit the broader applicability of our findings to bovine TB as a whole. Future studies should incorporate these virulent clinical strains to enhance the relevance of the results. Additionally, expanding the sample size to include more indigenous and crossbred cattle, as well as both TB-positive and TB-negative animals, will provide a more comprehensive understanding of susceptibility and resistance in cattle.

Applying a combination of genomic, transcriptomic, and proteomic approaches in future research will also help elucidate the global immune response signature associated with TB resistance in cattle, potentially leading to better disease management strategies.

In conclusion, our findings provide strong evidence that the superior immune response of Sahiwal cattle to *M. tuberculosis* is driven by distinct transcriptional and immunological mechanisms. The differential expression of key immune-related genes and pathways between indigenous and crossbred cattle underscores the importance of breed-specific immune profiling for understanding TB resistance. The transcriptional signature identified in this study offers valuable insights into potential biomarkers for disease resistance, with implications for developing targeted breeding strategies and interventions to combat TB in cattle. Further research into these molecular pathways could pave the way for novel host-directed therapies aimed at enhancing immune responses in susceptible breeds.

## Statements and Declarations

### Ethics Statement

All the experiments were reviewed and approved by the Institutional Biological Safety Committee (IBSC, Approval No. IBSC/2018/NIAB/BD/001) of the National Institute of Animal Biotechnology, Hyderabad, and animal experiments were approved by the Animal Ethics Committee of the West Bengal University of Animal and Fishery Sciences, Kolkata, India [Approval No. IAEC/22 (B), CPCSEA Reg. No.763/GO/Re/SL/03/CPCSEA), Committee for the Purpose of Control and Supervision on Experiments on Animals, India]. Blood collection from the jugular vein and tuberculin test on cattle were performed by a trained Veterinarian, and all the procedures were performed following relevant guidelines and regulations.

### Data Availability

The transcriptome data from this study have been made publicly available in the NCBI GEO database under accession number **GSE277021**. All other data have been provided as Supplementary data files and indicated in the manuscript.

### Software and Database

For pathway enrichment analysis and key gene selection, several publicly accessible databases were employed. Among the primary databases used were NCBI-GEO, g:profiler, AmiGO, and InnateDB. Detailed information on all databases and software utilized is available in **Supplementary Data File S6**.

### Author Contributions

The project was conceived and designed by BD. Experiments were performed by RK, SG, HKM, US, and BD. Data analysis was carried out by RK, SG, US, and BD. Contributed reagents, materials, analysis tools, and facilities: US and BD. The manuscript was written by RK, SG, and BD. All authors reviewed and edited the manuscript.

## Acknowledgments

We gratefully acknowledge Prof. Sharmistha Banerjee, coordinator of the University of Hyderabad – NIAB joint BSL3/ABSL3 facility at the University of Hyderabad, India. We thank the technical support staff of NIAB and UoH for their invaluable assistance in facilitating the BSL-3 and other laboratory-based experiments. We also thank the technical support staff of WBUAFS for their help during the experiments on cattle and sample collection.

## Supplementary information

Please see the supplementary information file.

