## Supplementary information for "Transcriptomic profiling identifies breed-specific immune signatures of Tuberculosis susceptibility in cattle"

**Title:**

**Affiliations:**

[#]Joint first author

***Corresponding Author:**

Dr. Bappaditya Dey

Scientist-F

National Institute of Animal Biotechnology

ORCID ID # 0000-0003-2728-4683

**Supplementary Fig. S1**

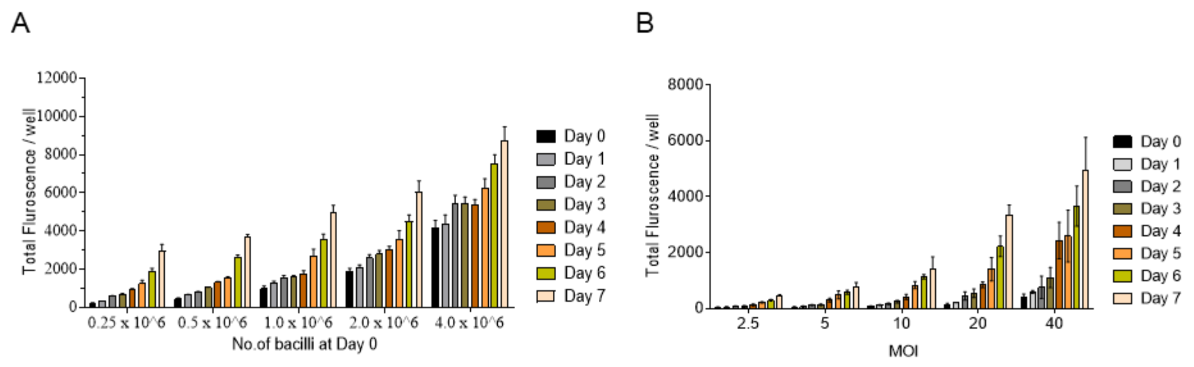


**Supplementary Fig. S1. Association of bacterial fluorescence and CFU.** *Mycobacterium tuberculosis* H37Rv expressing tdTomato fluorescence was generated. Assessment of the relationship of fluorescence with bacterial CFU was measured by two methods: (A) Fluorescent bacteria were inoculated into 7H9 broth at varying concentrations (0.25, 0.5, 1, 2, and 4 million CFU), and changes in fluorescence were monitored over 7 days. The total fluorescence intensity per well was plotted against bacterial concentrations on day 0. (B) BOMAC cells were infected with the reporter strain at different multiplicities of infection (MOI: 2.5, 5, 10, 20, and 40 bacteria per cell) and fluorescence intensity was tracked daily for 7 days post-infection. A strong positive correlation was observed between bacterial CFU and fluorescence intensity in both experiments. An MOI of 1:10 was selected for *ex vivo* mycobacterial growth assays using bovine PBMCs.

**Supplementary Fig. S2**

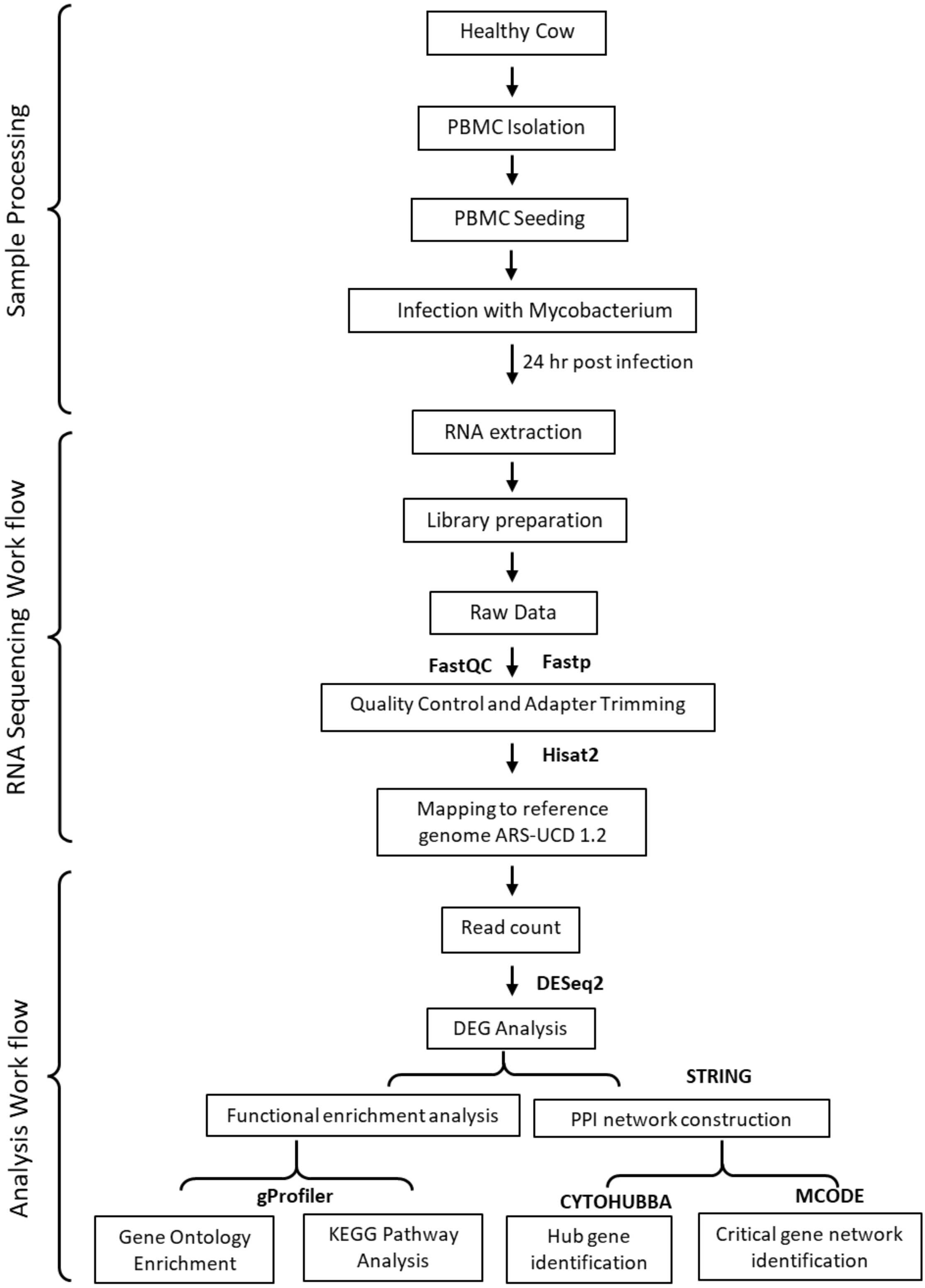


**Supplementary Fig. S2. Transcriptome sequencing and analysis workflow.**

### Supplementary Fig. S3

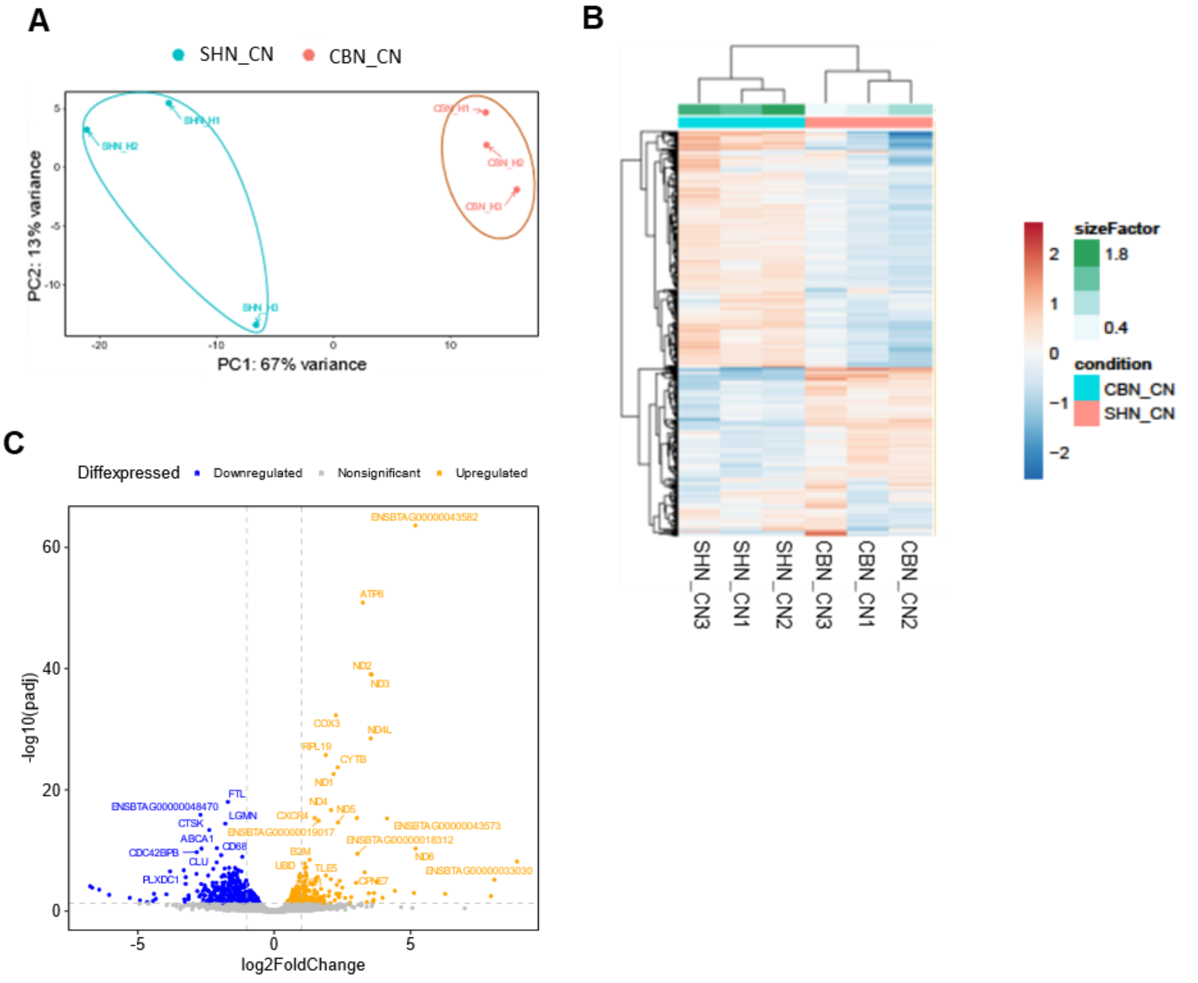


**Supplementary Fig. 3. Comparative transcriptome analysis of uninfected PBMCs from Sahiwal vs crossbred SHF cattle. (A)** The PCA plot representing separation between uninfected PBMC groups- Sahiwal vs. SHF; (**B**) Heatmap of expression profiles of top 500 genes displaying the relative expression levels of uninfected groups- Sahiwal vs. SHF; (F, G) Volcano plot depicting upregulated (orange) and downregulated (blue) differentially expressed genes (DEGs) in the uninfected- Sahiwal vs. SHF with an FDR < 0.05 and Log2FC > 1.

### Supplementary Fig. S4

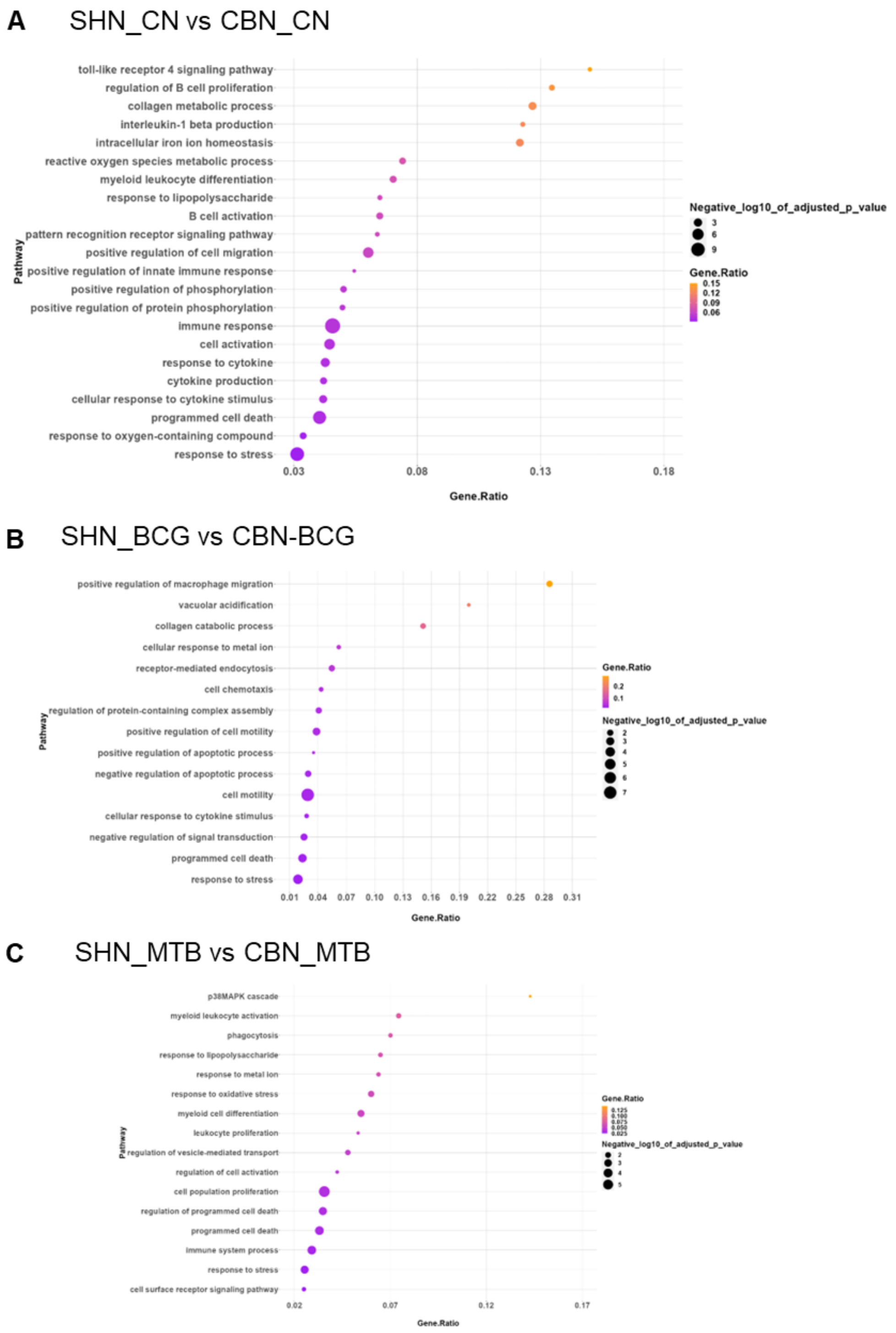


**Supplementary Fig. S4. Pathway enrichment analysis of the down-regulated genes.** Major biological pathway down-regulated in case of Sahiwal compared to the crossbred PBMCs, in different experimental conditions: (A) Uninfected, (B) *M. bovis BCG* infection, and (C) *M. tuberculosis H37Rv* infection. The comparison of DEGs was carried out between Sahiwal and crossbred cattle, using crossbred DEGs as the reference. Analysis was conducted using g:Profiler, and figures were generated with R-Studio.

**Supplementary Fig. S5**

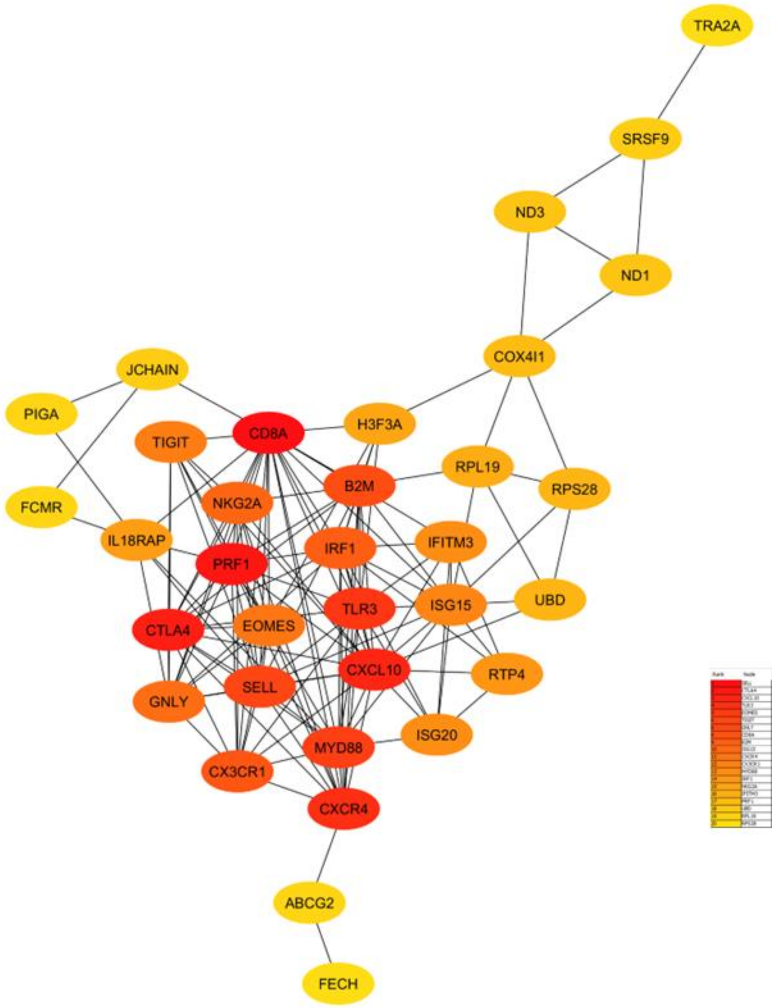


**Supplementary Fig. S5. Key hub nodes identified using the CytoHubba plugin in Cytoscape software.** The figure highlights the interconnected nodes within the global gene network based on the 34 shortlisted DEGs in *M. tuberculosis*-infected groups- Sahiwal vs crossbred, based on 7 different analysis features using CytoHubba and plotted by MCC method.

**Supplementary Table S1: Plasmids and mycobacterial strains used in the study**

| Name | Description | Source |
|---|---|---|
| **Plasmids** | | |
| pMSP12::mCherry | Mycobacterial reporter plasmid expressing mCherry | Addgene plasmid # 30169 |
| pTEC27-Hyg | Mycobacterial reporter plasmid expressing tdTomato | Addgene plasmid # 30182 |
| **Mycobacterial strains** | | |
| *M. tuberculosis H37Rv* | *M. tuberculosis* H37Rv strain- BSL3 grade virulent strain | Prof. S. Banerjee, University of Hyderabad, India |
| *M. bovis BCG* | *M. bovis BCG* Danish 1331 vaccine strain | Prof. S. Banerjee, University of Hyderabad, India |
| *M. tuberculosis* H*37Rv*- tdTomato | *M. tuberculosis* H37Rv strain- harbouring pTEC27-Hyg | This study |
| *M. bovis BCG* - mCherry | *M. bovis BCG Danish 1331* strain harboring pMSP12::mCherry | *Kumar et al. 2023 (ref,18)* |

### Supplementary Table S2: qRT- PCR primers

| Primer Name | Nucleotide Sequence | Gene description |
|---|---|---|
| *bISG15 | F, 5′-tgtgccaaaaggagcgtgta-3′<br>R, 5′-cccaccccgaagacgtagat-3′ | Interferon stimulated gene 15 |
| bEOMES | F, 5′-acaactatgattcatcccatcaga-3′<br>R, 5′-ggcacggttctctcaccatt-3′ | Eomesodermin |
| bCTLA4 | F, 5’-gccctgcactgccctatttt-3′<br>R, 5′-cgtcagctttgcctgaagac-3′ | Cytotoxic T-lymphocyte associated protein 4 |
| bSELL | F, 5′-tcaccccсttgtcgactttg-3′<br>R, 5′-gggctggaccaatttccaga-3′ | L-selectin |
| bTLR3 | F, 5′-tcgtcacagtaagtgaaggcaa-3′<br>R, 5′-acagtacatttgtgggcagaag-3′ | Toll-like Receptor 3 |
| bMYD88 | F, 5′-ggcagctggaacagacaaac-3′<br>R, 5′-cgacggcacctcttctcaat-3′ | Myeloid differentiation primary response gene 88 |
| bIRF1 | F, 5′-agcttacctttgggatgagtgg-3′<br>R, 5′-agccaagggcaatgactgtat-3′ | Interferon regulatory factor 1 |
| bCXCL10 | F, 5’-tcctcgaacacggaaagagg-3’<br>R, 5’-gtccacggacaattagggct-3’ | C-X-C motif chemokine ligand 10 |
| bRPLP0 | F, 5′-cttcattgtgggagcagaca-3′<br>R, 5′-ggcaacagtttctccagagc-3′ | 60S acidic ribosomal protein large |
| *b, Bovine. | | |
